# Dissecting neurofilament tail sequence-phosphorylation-structure relationships with multicomponent reconstituted protein brushes

**DOI:** 10.1101/2024.05.02.592230

**Authors:** Erika A. Ding, Takashi J. Yokokura, Rui Wang, Sanjay Kumar

## Abstract

Neurofilaments (NFs) are multi-subunit, bottlebrush-shaped intermediate filaments abundant in the axonal cytoskeleton, with “bristles” composed of the subunits’ disordered tail domains. Precisely how the tails’ variable charge patterns and repetitive phosphorylation sites mediate their conformation within the brush remains an open question in axonal biology. We address this problem by grafting recombinant NF tail protein constructs (NFL, NFM, and NFH) to functionalized substrates, forming phosphorylatable brushes of defined stoichiometry. Atomic force microscopy reveals that NFM-based brushes are highly extended, while brushes incorporating the much larger NFH are surprisingly compact even after multisite phosphorylation. A self-consistent field theory predicts multilayered brush morphologies for NFM and phosphorylated NFH brushes. Further experiments with designed mutants reveal that N-terminal negative charges in NFH repel phosphorylated residues to generate the multilayer morphology and that charge segregation in NFM promotes collapsed conformations, lending new insight into how NF tail sequence features determine protein brush conformation.

## Introduction

Neurofilaments (NFs) are bottlebrush-shaped, neuron-specific intermediate filaments that fill the axonal cytoplasm and contribute to axonal caliber^1^. NFs are composed of multiple protein subunits with unique C-terminal tail domains which protrude from the filament core, forming a layer of protein surrounding each filament (Fig. 1A-B)^2,3^. Being intrinsically disordered regions (IDRs), these tail domains do not take on well-folded structures but rather sample a conformational ensemble^4^. This layer can therefore be viewed as a polymer brush, with the tail domain proteins as polymers tethered to a nanoscale cylindrical support. The conformations adopted by polymers in a brush reflect contributions from the elastic energy of polymer deformation, interactions between monomers and solvent, and electrostatic interactions between charged monomers and dissolved ions. In the context of the NF tail domains, these interactions are governed by the subunits’ amino acid sequences.

**Figure 1.**
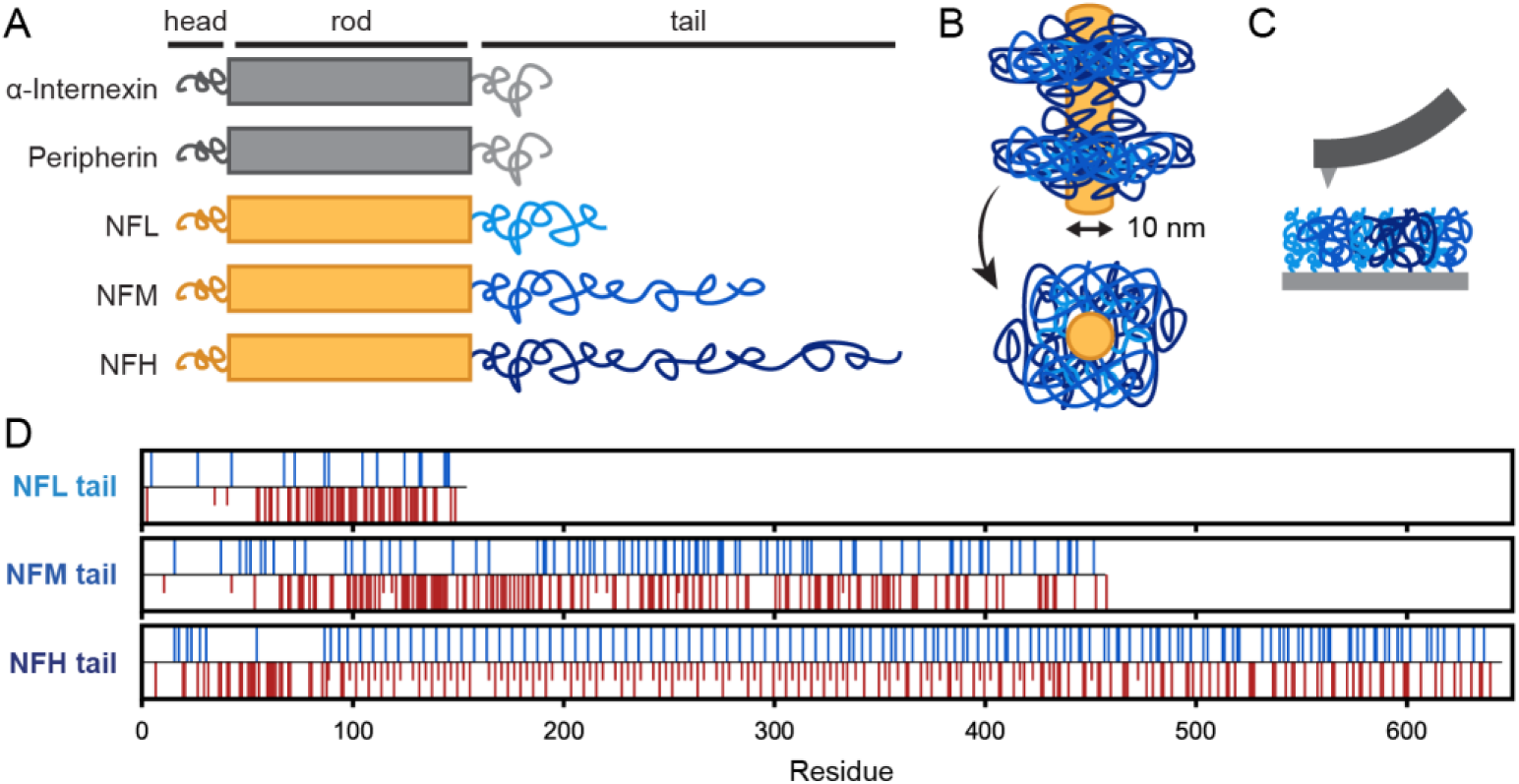
NF tails with unique sequence features form a mixed brush. (A) Diagrams of the neurofilament protein domains. (B) Schematic of assembled neurofilament with surrounding tail domain layer. (C) Schematic of in vitro brush system and AFM measurement. (D) Charge properties of NF tail constructs at the neutral pH used in this work. Blue bars indicate lysine or arginine residues, red bars indicate glutamate or aspartate residues, and half-length red bars indicate phosphorylated serines or threonines as detected by peptide LC-MS.

*In vivo*, the NF tail domains have been proposed to mediate inter-filament interactions and crosslinking ^5–7^, to act as space-filling buffers between filaments^8,9^, and possibly to support the mechanical properties of the axons where NFs are found^10,11^, with these functions disrupted by NF aggregation in many neurodegenerative conditions^3,12^. However, the field still lacks a sequence-to-structure paradigm linking NF tail amino acid sequence features to the structure and organization of the NF tail layer.

Efforts to define sequence-structure relationships are complicated by the heteropolymeric nature of native NFs, which are always formed from more than one subunit protein. The three best-studied NF subunits are NF-Light, -Medium, and -Heavy (NFL, NFM, and NFH), and the tail domains of these subunits are highly charged and variable in length. Neurons tune NF function not only by differentially regulating the subunits at transcriptional and post-transcriptional levels^13^, but also by multisite phosphorylation, especially at repetitive lysine-serine-proline (KSP) motifs targeted by various proline-directed kinases. The availability of these phosphorylation motifs varies by subunit; for example, the rat NFM tail sequence contains 5 KSP motifs, while the rat NFH tail contains 52 such motifs^2,14^. Multisite phosphorylation brings additional negative charge to the sequence, changing the monomer chemistry within the protein brush. However, the effect of multisite phosphorylation on protein conformation remains unclear.

Various approaches have been implemented to understand the roles of the different NF tail domains in governing NF brush morphologies. Studies of reconstituted NF hydrogels have suggested that NFM decreases inter-filament spacing, while NFH supports expanded inter-filament spacing especially after phosphorylation^11,15^. However, in mouse genetic models, deletion of the NFM tail decreases NF spacing^16^, while deletion of the NFH tail has no effect^17^. Both hydrogel and mouse genetic models infer NF brush morphology via inter-NF spacing, which cannot separate the effects of NF tail conformation from inter-filament interactions. Protein conformation can be predicted by coarse-grained computational approaches, which have suggested brush morphologies with NFM extending to the periphery of the brush and NFH doing so after phosphorylation^18,19^. However, these protein conformation predictions cannot be experimentally corroborated in complex hydrogel or animal models.

Here we employ a reductionist, well-controlled experimental system combined with a self-consistent field theory to directly probe ensemble-average conformation within a protein brush. With these approaches we explore how brush morphology is governed by sequence-encoded biochemical features such as charge pattern, chain length, and multiple phosphorylation sites in single- and multi-component NF tail brushes.

## Results

### NF tail constructs form ternary protein brushes of controlled density, composition, and phosphorylation state

Building from a previously developed platform^20^, we grafted NF tail proteins to a maleimide-functionalized glass or silicon surface (Fig. 1C). This system recapitulates the surface-tethered context of the NF tails *in vivo*, while also allowing control over the grafting density and composition of mixed brushes made of multiple subunits. Protein constructs based on *R. norvegicus* NFL, NFM, and NFH tail domains were recombinantly produced in *E. coli*, allowing sequence modifications and the introduction of a cysteine residue near the N-terminus for end-directed covalent grafting via a thiol-maleimide Michael addition (Supporting Fig. 1). The purified proteins, which like other highly charged IDPs migrate anomalously in SDS-PAGE^21,22^, were identified by LC-MS (Supporting Fig. 2). These constructs can be phosphorylated at KSP motifs by purified recombinant kinase ERK2 either in solution or after surface grafting ^23^. LC-MS analysis of trypsin-digested phosphorylated proteins revealed 53 phosphorylated residues in the NFH tail construct used here, 8 residues in NFM, and 2 residues in NFL (Fig. 1D).

After allowing each protein to graft to a functionalized surface, we measured the density of grafted protein by spectroscopic ellipsometry. The grafting densities σ of single-component NFL, NFM, or NFH tail brushes were similar to one another (Fig. 2A), and similar to the density of 0.02 chains nm^-2^ previously reported using this grafting chemistry for an NFH tail construct^20^. These grafting densities are sparse compared to those of typical synthetic polymer brushes, which is unsurprising given the grafting-to reaction scheme and the long protein chains (molecular weights of NFL, NFM, and NFH tails are 17, 51, and 69 kDa, respectively). However, the present grafting densities are similar to the density of tail domains in a native NF. Assuming 32 subunits incorporated per cylindrical unit of diameter 10 nm and length 43 nm^24^, a native NF grafting density might be estimated as 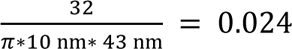.

**Figure 2.**
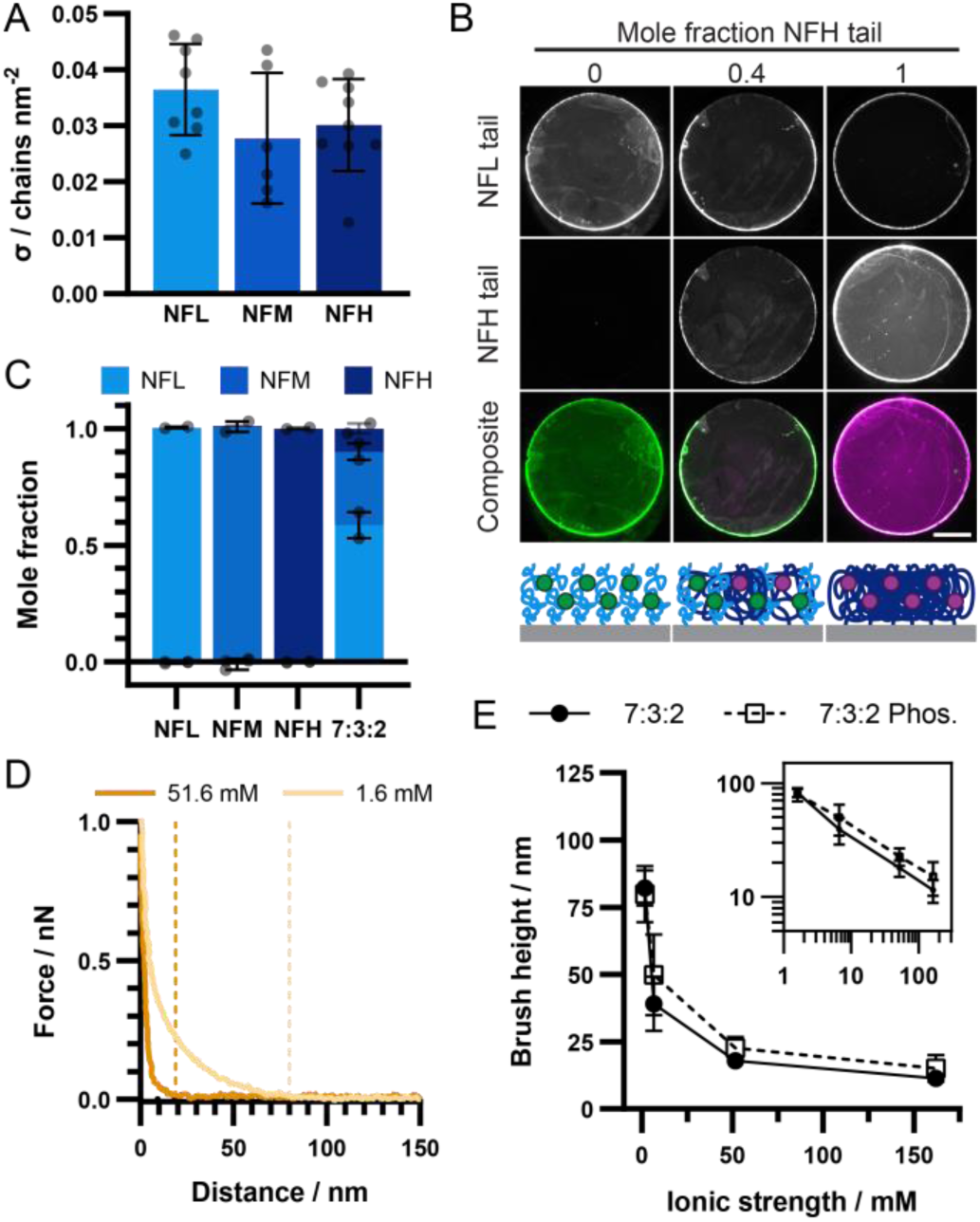
Ternary NF brushes of controlled grafting density and composition collapse with increasing solution ionic strength. (A) Grafting density of single-component brushes. Mean and standard deviation, n = 6-9 samples. (B) Representative images (top) and schematic (bottom) of NFL:NFH fluorescently labeled mixed brush, scale bar 5 mm. (C) Molar compositions of single-component brushes and 7:3:2 ternary brush. Bars show mean +/- range from each channel, n = 2 samples. (D) Representative curves from AFM brush height measurement at high (dark orange) and low (light orange) ionic strength. Dashed lines indicate extracted brush heights. (E) Collapse of ternary brush height with increasing ionic strength with (open squares) or without (closed circles) phosphorylation. Points show mean and standard deviation, n=5 samples. Inset: log scale view.

To produce multicomponent brushes, we allowed a mixture of proteins in solution to simultaneously co-graft to the surface. We empirically related the composition of the protein mixture applied to the surface and the brush molar composition using relative fluorescence. Proteins were lightly pre-labeled with amine-reactive fluorophores, then allowed to co-graft on the surface and visualized with a laser scanner platform (Fig. 2B, Supporting Fig. 3). By comparing fluorescence intensity from mixed brushes to the intensity from a single-component reference sample of known grafting density, we calculated the grafting density of each component in the mixed brush, and the resulting molar brush composition (Methods).

*In vivo*, NFs are found in a range of protein compositions that depend on many factors^25–27^. However, one commonly described stoichiometry is a molar NFL:NFM:NFH ratio of 7:3:2 ^28^, so we chose a protein mixture that resulted in a ternary brush composition close to this ratio (Fig. 2C). We then measured brush height with atomic force microscopy (AFM) under a variety of solution conditions. This force-based metric records the position at which the AFM probe first detects resistance as it approaches the sample, relative to the position of near-vertical force response where the probe is in contact with the underlying glass substrate. Representative force-distance curves demonstrate the difference in brush height for a 7:3:2 ternary brush at low (1.6 mM) or high (51.6 mM) solution ionic strength, showing that the brush is more extended at low ionic strength (Fig. 2D). All brush height measurements in this work were performed at pH 7.4 to reflect physiological pH, and all samples used for brush height measurements were made with unlabeled proteins.

The height of ternary NF tail brushes decreased with increasing solution ionic strength according to a power law with exponent -0.4176 (Fig. 2E). This collapse is more dramatic than would be expected for a classic “salted” polyelectrolyte brush, which exhibits a scaling law exponent of -0.33 with ionic strength^29^. We note that the brush height of ∼15 nm near physiological ionic strength is consistent with the range inferred from electron micrographs of mouse axons, where an observed 30-50 nm inter-NF spacing would correspond to a 10-nm filament core with two interposed brushes of 10-20 nm^17,30^. *In vivo*, extensive phosphorylation of NFM and NFH tail domains has been proposed to contribute to axonal NF organization and function. Therefore, we phosphorylated ternary NF tail brushes *in situ*, and observed slightly expanded brush heights and a concomitant decrease in magnitude of the power law exponent (Fig. 2E, Supporting Table 1).

### Protein composition, phosphorylation, and ionic strength govern expansion of binary brush systems

As the NFM and NFH tails are both long and highly charged chains, we particularly wanted to isolate the contributions of these subunits to brush morphology. Because NFL is required for filament assembly *in vivo*^1^ and would therefore always be present in the native NF tail brush, we varied the mole fraction of NFM or NFH tails against a background of NFL in a series of binary mixed brushes with constant total protein grafting density. We again combined spectroscopic ellipsometry and relative fluorescence to calculate f_NFM_ or f_NFH_, the mole fraction of NFM or NFH within a given brush. We empirically correlated the composition of the applied protein mixture and the resulting brush composition (Fig. 3A). NFL:NFM binary brush compositions were fit using a quadratic function resulting in a standard deviation of the residuals (Sy.x) of 0.11, while NFL:NFH brush compositions were linearly correlated to grafting solution composition with Sy.x 0.093. Total protein grafting density on the surface showed no trend with brush composition and was characterized by a standard deviation of 0.012 chains nm^-2^ over NFL:NFM samples and 0.0085 chains nm^-2^ over NFL:NFH samples (Supporting Fig. 4).

**Figure 3.**
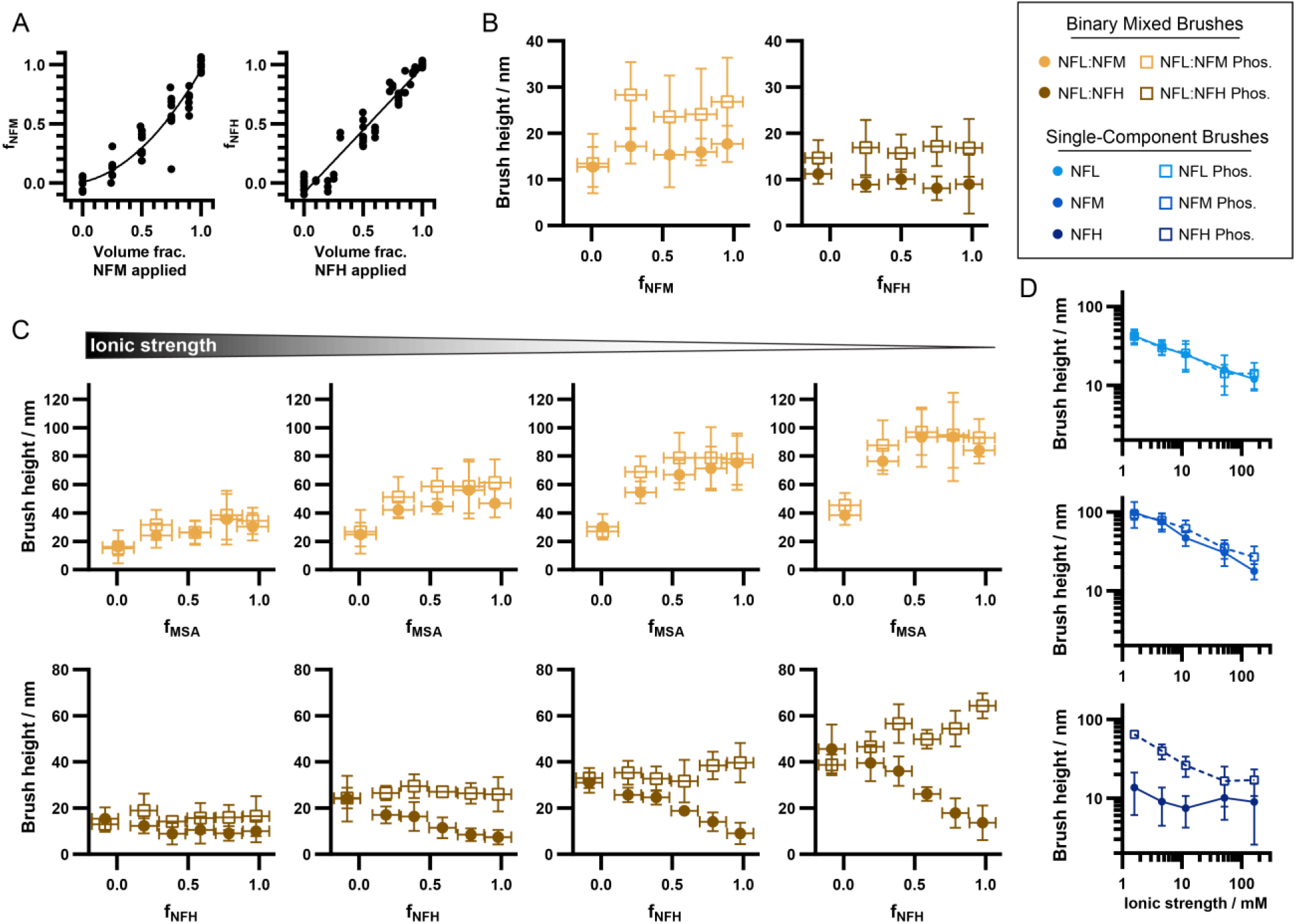
NFM contributes to brush expansion while the effect of NFH is phosphorylation-dependent in binary mixed brushes. (A) Brush compositions for binary mixtures of NFL:NFM (left) and NFL:NFH (right) as a function of protein mixture applied to the surface. Empirical fits (solid lines) allow control of brush composition within sample error. Points represent individual samples. (B) Heights of NFL:NFM (left) and NFL:NFH (right) mixed brushes in phosphate-buffered saline with (open squares) or without (closed circles) phosphorylation. Points represent mean and standard deviation, n = 4-8 samples. X-error bars show standard deviation of the residuals from the relevant fitting in (A). (C) Heights of NFL-NFM (top row) and NFL-NFH (bottom row) mixed brushes in lower ionic strength solutions with (open squares) or without (closed circles) phosphorylation. Left to right: 51.6, 11.6, 4.6, 1.6 mM ionic strength. Points represent mean and standard deviation, n = 4-8 samples. X-error bars show standard deviation of the residuals from the relevant fitting in (A). (D) Single-component brush heights against solution ionic strength for NFL (top), NFM (middle), or NFH (bottom) tail construct brushes. Points represent mean and standard deviation, n = 4-8 samples.

We measured the height of a series of NFL:NFM or NFL:NFH mixed brushes in phosphate-buffered saline (Fig. 3B). Against a background of NFL, NFM tended to increase the brush height, with phosphorylation amplifying this effect. Meanwhile, introducing NFH only minimally affected brush height, though phosphorylation slightly expanded brushes incorporating NFH.

To investigate the role of charge interactions in binary mixed brushes, we tuned ionic strength with NaCl in a weakly buffered system. At lower ionic strengths, all brushes were more expanded and the effects of protein composition were more pronounced, highlighting the importance of electrostatic effects in this system (Fig. 3C). Binary brush heights were generally monotonically dependent on composition, though this dependence was nonlinear for NFL:NFM mixed brushes. Increasing the fraction of NFM resulted in more expanded brushes with only slight changes upon phosphorylation. Conversely, while phosphorylated NFH tended to increase the brush height, non-phosphorylated NFH resulted in more collapsed brushes.

To dissect these effects more finely, we examined single-component NFL, NFM, or NFH brushes. Among these three, NFM brushes were the most extended at all ionic strengths measured (Fig. 3D). As expected, NFL produced shorter brushes than NFM. Intriguingly, NFM tails were able to form more expanded brushes than phosphorylated NFH at all ionic strengths, despite the NFH sequence being significantly longer than that of NFM and being expected to contain more net negative charge than NFM after phosphorylation. Increasing the concentration of ATP did not further expand NFH brushes, suggesting saturation of available phosphosites (Supporting Fig. 5). We note that NFL brush heights do not change upon phosphorylation, while NFM brushes expand slightly at higher ionic strengths. NFL and NFM single-component brushes also follow a power law within the measured ionic strength regime irrespective of phosphorylation state (Supporting Table 1). However, non-phosphorylated NFH brushes take a collapsed morphology even at low ionic strength, and phosphorylated NFH brushes collapse below a solution ionic strength of ∼50 mM before plateauing at higher ionic strengths.

### SCFT predicts sequence-dependent multilayer brush morphologies

To explore chain organization within protein brushes, we turned to self-consistent field theory (SCFT). SCFT is well-suited to modeling inhomogeneous polymer systems, providing polymer density profiles that are experimentally inaccessible. We applied a continuous-space SCFT previously developed for NF tail brushes^31^, which systematically includes the elastic energy due to protein chain deformation, short-range residue hydrophobicity via the Flory-Huggins parameter, and electrostatic interactions. We employed a coarse-graining procedure in which each amino acid sequence is mapped onto a multi-block copolymer model, where each block is parameterized by the average hydrophobicity and charge of its constituent residues at the chosen pH. We applied this model to single-component NF construct brushes. Based on the best fit with the experimental data (Fig. 4A-C, insets), the same Kuhn length *b* and segment volume *v* were chosen for all constructs except for phosphorylated NFH, where *v* was increased to account for the volume change due to the phosphates (Supporting Note 1).

**Figure 4.**
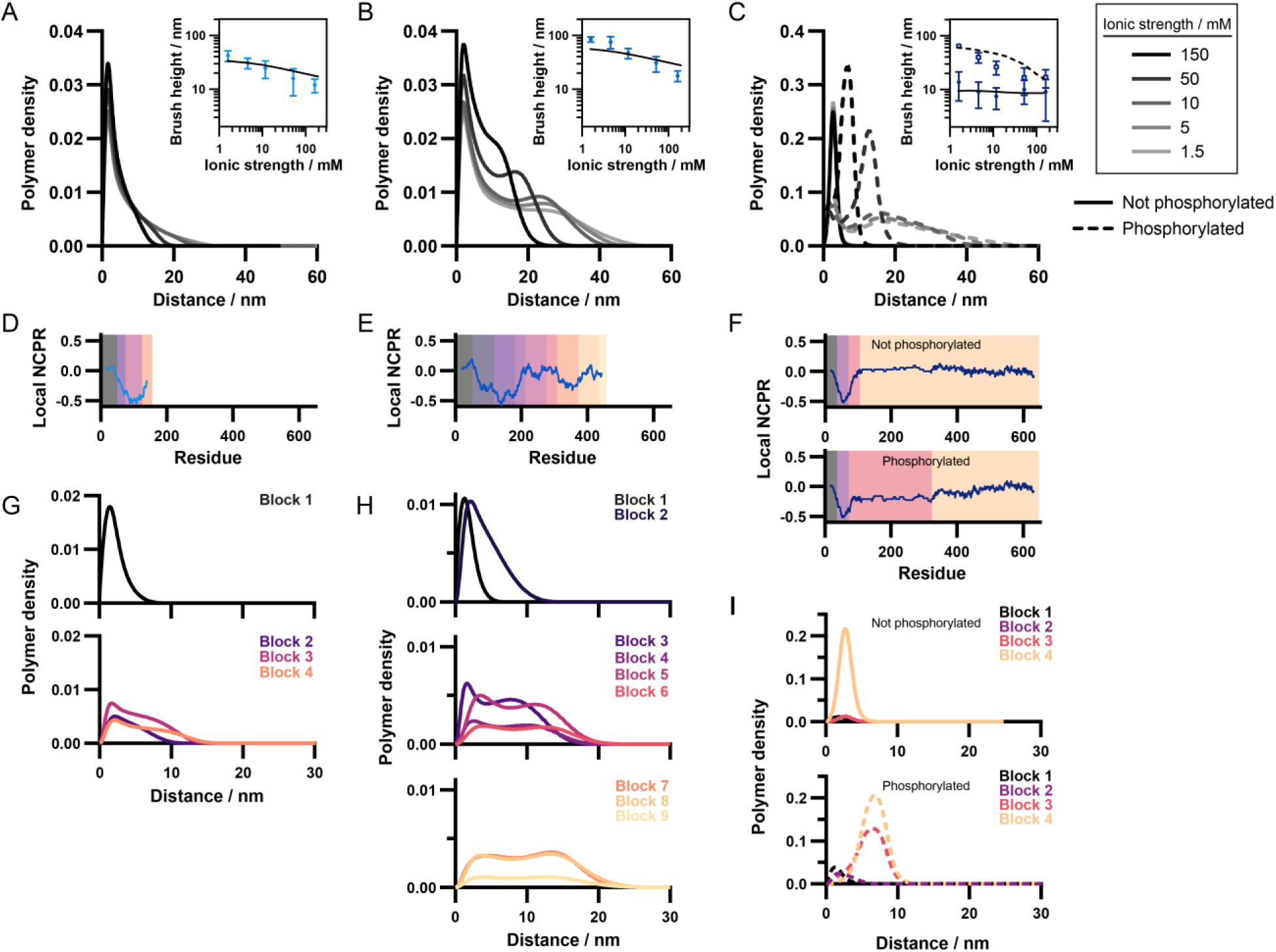
SCFT predicts multilayer sub-brush structures. (A,B,C) Calculated polymer density profiles of NFL (A), NFM (B), or NFH (C) single-component brushes showing collapse with ionic strength. NFH polymer density profiles are shown with (dashed lines) or without (solid lines) phosphorylation. Insets: predicted brush heights (lines) and experimental data (points; mean and standard deviation); NFH is shown with (open squares data, dashed line fit) or without (closed circles data, solid line fit) phosphorylation. (D,E,F) Local net charge per residue for NFL (D), NFM (E), non-phosphorylated NFH (F, top), or phosphorylated NFH (F, bottom). Colors indicate charge blocks used in the model. (G,H,I) Polymer densities separated by charge block for NFL (G), NFM (H), or NFH (I) brushes at 150 mM ionic strength. NFH profiles show phosphorylated (dashed) and nonphosphorylated (solid) conditions.

Polymer density distributions calculated from the SCFT predict that NFL brushes take a single-layer morphology which slightly condenses with increasing ionic strength (Fig. 4A). By contrast, NFM brush polymer densities exhibit a peak near the surface and a shoulder further away (Fig. 4B), indicating the coexistence of a condensed and a more diffuse layer. As ionic strength increases, the shoulder in the polymer density profile merges into the peak, indicating that outer diffuse layer collapses towards the surface. For NFH, brush morphology depends strongly on phosphorylation state. While non-phosphorylated NFH brushes form a fully condensed layer, phosphorylated NFH brushes are predicted to consist of a relatively small inner layer and an expanded diffuse layer, which collapses with increasing ionic strength (Fig. 4C).

Each protein is a polyampholyte with sequence-defined charge patterns. These patterns can be described by a moving-window local net charge per residue (NCPR) and were used to divide each sequence into blocks of similar charge. While NFL contains a relatively neutral block followed by three blocks of net negative charge (Fig. 4D, blocks denoted by shading), NFM contains a charge distribution with two negative peaks and is segregated into nine blocks (Fig. 4E). Four blocks are needed to model NFH, including a notable N-terminal block of dense negative charge (Fig. 4F, top). Upon phosphorylation, changes to the local NCPR profiles in NFL and NFM are negligible (data not shown), but the central section of NFH becomes more negatively charged (Fig. 4F, bottom). Of the four blocks modeling phosphorylated NFH, Block 3 contains the majority of the phosphorylation sites.

To understand how charge patterning affects the brush morphology, we examine the polymer density distributions of each subunit’s constituent blocks at a high ionic strength of 150 mM (Fig. 4 G-I). For NFL, while the relatively charge-neutral N-terminal region (Block 1) is predicted to remain close to the surface, the more negatively charged C-terminal blocks (Blocks 2-4) extend further into the solution (Fig. 4G). In NFM, while Blocks 1 and 2 mostly populate the condensed layer, Blocks 3-9 have a bimodal distribution existing in both the condensed and the diffuse layer (Fig. 4H). This distribution suggests that the dense negative charges in Blocks 3 and 7 favor stretched conformations in these blocks due to local charge repulsion, causing adjacent blocks to be dragged to the outer layer due to chain connectivity. For unphosphorylated NFH, the N-terminal negative charges are almost completely neutralized by the positive charges carried by the remaining residues. The brush behaves effectively as a charge neutral polyampholyte and is insensitive to changes in ionic strength, with all constituent blocks condensed to a single layer due to backbone hydrophobicity (Fig. 4I, top). For phosphorylated NFH, the surface-adjacent inner layer is predicted to be composed of Blocks 1 and 2, while the outer layer is predicted to be composed of Blocks 3 and 4 (Fig. 4I, bottom). The highly negatively-charged Block 2 repels negatively charged phosphates in Blocks 3 and 4, preventing them from condensing to the surface.

### Sequence-encoded charge patterns in NFM and NFH govern brush morphologies

The SCFT results predict that chain organization within NF tail brushes depends strongly on sequence charge distribution. We therefore returned to our recombinant protein system to probe how specific charge patterns in the NFM and NFH tails govern brush extension. In NFH, we focused on charged regions other than the extensively phosphorylated KSP repeat region. We first deleted the dense block of negative charge near the N-terminus of the protein, forming the construct NFH Δ36-72 (Fig. 5A, Supporting Fig. 1). We confirmed the identity of the purified protein by LC-MS (Supporting Fig. 2) and validated brush formation by spectroscopic ellipsometry, then measured the response of the resulting brush to solution ionic strength (Fig. 5B). Although we achieved lower brush densities than for WT NFH (Supporting Fig. 6), we found that without phosphorylation, the NFH Δ36-72 construct formed a somewhat more expanded brush than the wild type construct, especially at low ionic strengths. This finding suggests that residues 36-72 and other negative charges near the grafting point may help attract the slightly positively charged KSP repeat region into a compact brush. Upon phosphorylation, the NFH Δ36-72 brush heights were only slightly more compact than the wild type sequence, and the power-law collapse regime of phosphorylated NFH Δ36-72 brush height began to plateau around 10 mM ionic strength, a lower threshold than for the WT NFH brush. At higher ionic strengths, the SCFT predicts that the multilayer expulsion of the phosphorylated KSP repeats that was observed in the WT brush is no longer favorable (Fig. 5D), though a thin dilute layer is still observed at ionic strengths below 10 mM (Supporting Fig. 7).

**Figure 5.**
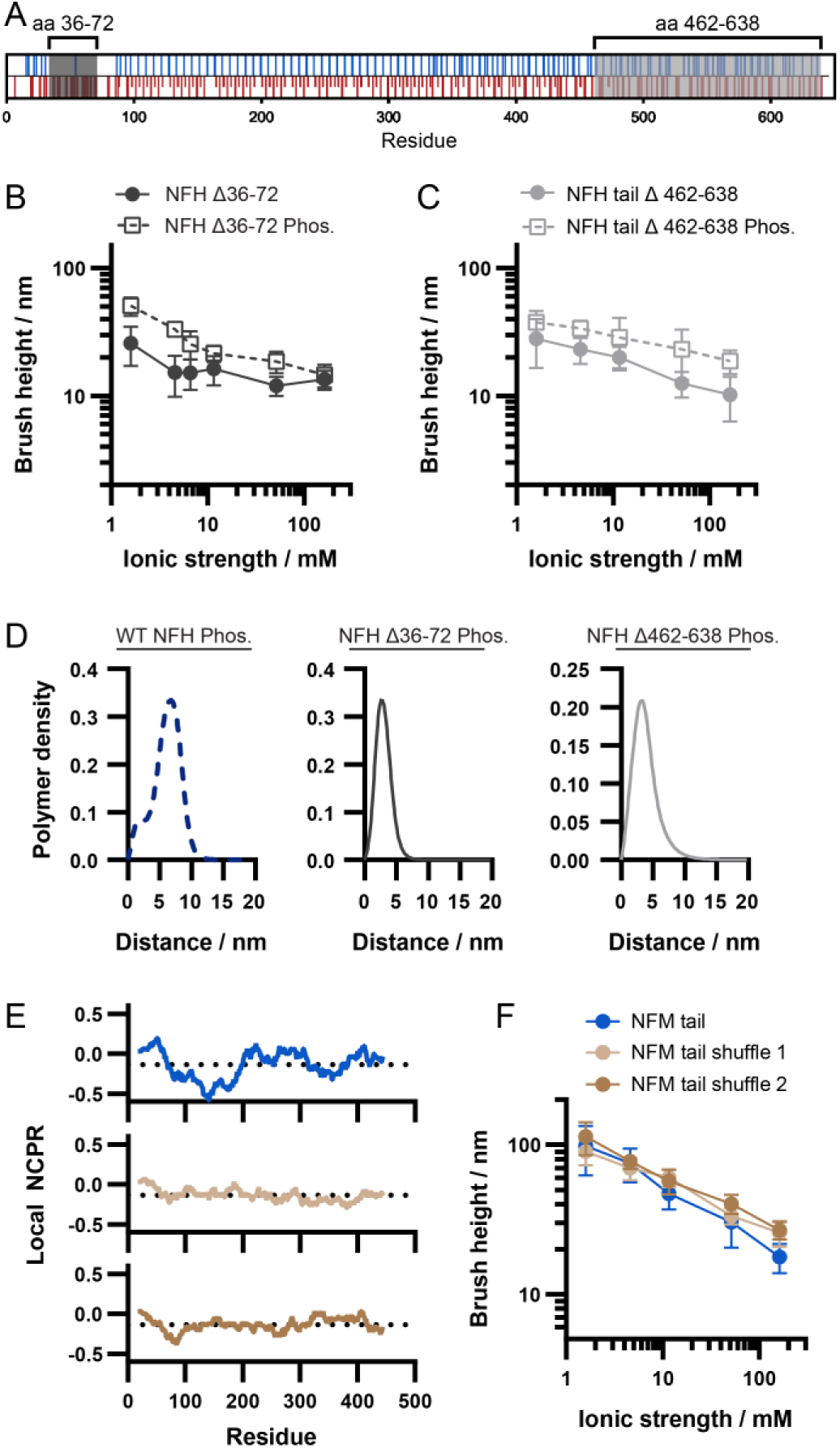
Charge patterns in NFM and NFH govern brush expansion. (A) Sequence charge diagram indicating the regions of NFH targeted in deletion constructs. (B) Experimentally measured heights of NFH tail Δ36-72 brushes (mean and standard deviation, n = 6-8 samples) with (open squares) and without (closed circles) phosphorylation. (C) Heights of NFH tail Δ462-638 single-component brushes with (open squares) or without (closed circles) phosphorylation. Mean and standard deviation, n = 9-14 samples. (D) Predicted polymer density profiles for phosphorylated NFH-derived brushes comparing wild type and deletion constructs at 150 mM ionic strength. (E) Local net charge per residue for wild type NFM tail construct (top) and two charge-shuffled variants (middle, bottom). Dotted line indicates average charge over the entire sequence. (F) Heights of NFM charge shuffled variant brushes compared to wild-type NFM. Mean and standard deviation, n = 6-8 samples.

We also wanted to know whether the C-terminal relatively charge-neutral block of NFH was contributing to brush height. We deleted this block to form the construct NFH Δ462-638 (Fig. 5A, Supporting Fig. 1, Supporting Fig. 2, Supporting Fig. 6) and measured the brush height in response to ionic strength (Fig. 5C). Interestingly, the NFH Δ462-638 brush height did not plateau in the ionic strength regime measured here, and resulted in a much lower-magnitude power law exponent than other brushes measured (Table S1). However, near physiological ionic strength the NFH tail Δ462-638 brush heights were very close to the WT NFH brush irrespective of phosphorylation state. SCFT predictions of NFH Δ462-638 brush morphologies suggest that repulsion of the phosphorylated residues into an outer layer still occurs at lower ionic strengths, but near physiological ionic strength the layers merge and the brush occupies only one layer (Fig. 5D, Supporting Fig. 7).

Compared to NFH, the NFM tail sequence is relatively charge-segregated, with blocks of dense negative charge spaced by more neutral regions. One metric to quantify sequence charge segregation is κ, a parameter which is close to 1 for charge-segregated sequences and close to 0 for well-mixed sequences^32^. The wild type NFM tail sequence is characterized by κ = 0.15. We created two charge-shuffled mutants with more evenly spaced charges by rearranging the charged residues so that they alternated along the sequence, randomly distributing the excess negatively charged residues, to create the constructs NFM shuffle 1 and NFM shuffle 2 with κ of 0.05 and 0.06, respectively (Fig 5E, Supporting Fig. 1, Supporting Fig. 2, Supporting Fig. 6). The charge-shuffled NFM variants resulted in similar brush heights to the wild type protein at low ionic strengths, but were more able to resist brush collapse at high ionic strength (Fig. 5F) and exhibited slightly lower-magnitude power law exponents compared to the wild type sequence (Table S1).

## Discussion

Despite the abundance of NFs in the axonal cytoskeleton and the importance of the tail domains in governing NF architecture, our mechanistic understanding of how amino acid sequences determine NF tail protein conformation remains incomplete. Previous experimental studies of the NF tail domains have inferred brush size from axonal inter-NF spacing^17,33^, but these approaches do not readily allow manipulation of tail sequences, control of subunit stoichiometry, or direct measurement of brush height. Here, we combined a recombinant NF subunit platform and SCFT to directly probe how sequence- and phosphorylation-defined charge features govern protein conformation in single-component, binary, and ternary protein brushes. We find that protein composition regulates the effect of phosphorylation on brush height, with NFM expanding mixed brushes irrespective of phosphorylation state and multisite phosphorylation in NFH causing a conformational shift via repulsion against a region of N-terminal negative charge (Fig. 6).

**Figure 6.**
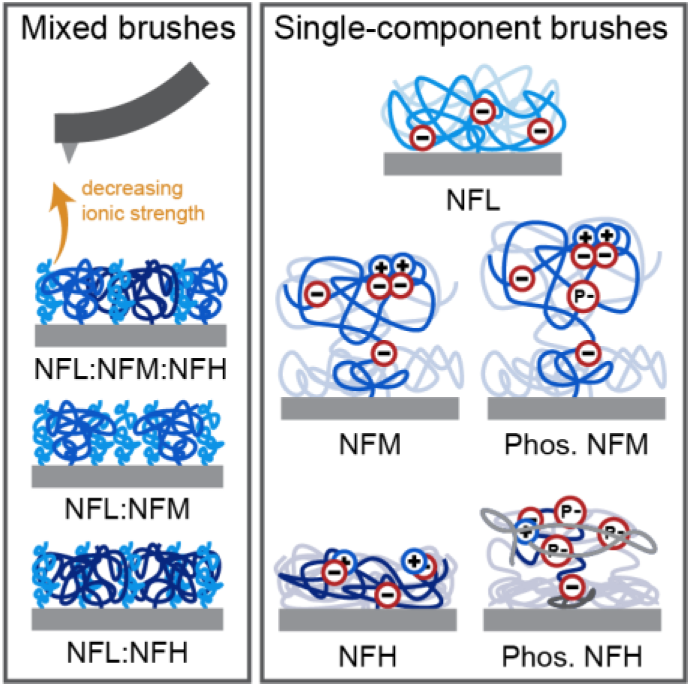
Summary. (left) Grafting recombinant proteins onto surfaces allows AFM-based height measurements of ternary, binary, and single-component brushes. All brushes swell with decreasing ionic strength. (right) Experimental and SCFT results indicate that near neutral pH and physiological ionic strength, NFL brushes form a single layer due to evenly spaced negative charge repulsion (top), while NFM brushes take on a condensed layer and a dilute layer that expands slightly after phosphorylation (middle). NFH is nearly net neutral before phosphorylation leading to a collapsed brush, while after phosphorylation an outer layer is formed by repulsion of the phosphates against an N-terminal negatively charged region, while the most C-terminal region does not further contribute to brush height (bottom).

The brush height data support a role for NFM in expanding the NF tail brush via repulsion between the segregated regions of dense negative charge, which causes a fraction of the protein chains to form an extended dilute layer. Interestingly, the brush height increases nonlinearly with NFM mole fraction in binary NFL:NFM brushes. Adding even a modest amount of NFM substantially increased the measured height of the brush to nearly the level of the pure NFM brush height (Fig. 3C), suggesting that the negative charge provided by the background NFL is enough to expel the dense negative charge blocks of NFM to form an outer layer.

Our study introduces novel charge-shuffled variants of the full-length NFM tail sequence, which enabled us to investigate the effects of charge distribution on brush height. In a single-component brush, more evenly shuffling the NFM sequence charges caused the brush to expand further. This finding in the context of surface-tethered IDPs complements evidence from IDPs free in solution^32,34–36^ and even specifically NFM tail fragments in solution^37^, where sequence charge desegregation tends to cause chain expansion. We observe that the charge shuffle effect on NFM tail brush height is negligible at low ionic strengths, possibly due to the lack of screening of significant charge repulsion between distant residues in the net-negative NFM sequence. Notably, the effect of charge patterning on IDP conformation has been suggested to be proline-dependent ^37,38^, so charge patterning may exert a different effect in NFH, which contains well-mixed charges and many relatively evenly distributed prolines (Supporting Fig. 1).

Our data also provide new perspective on previous conflicting findings regarding the effect of NFM tail phosphorylation. We find that the NFM tail expands only minimally upon phosphorylation in both binary and single-component brushes, in agreement with previous findings from Monte Carlo simulation^19,39^. In contrast, in reconstituted NF hydrogels NFM phosphorylation was observed to decrease the estimated NF tail brush thickness^11^. Inter-NF spacing is slightly smaller in phosphoincompetent NFM tail mice than in WT mice at 2 months of age, though this difference equalizes by 6 months of age^30^. Inter-NF spacing in this complex animal model may also reflect age-dependent compensatory phosphorylation effects, but the modest impact of NFM tail phosphorylation is consistent with our finding that this phosphorylation only minimally impacts brush height. Discrepancies between simulation, hydrogel, and mouse models may derive from the species-dependent availability of phosphorylation sites in the NFM tail sequences. NF hydrogel models^11^ have used native NFs purified from bovine spinal cord, where the NFM tail sequence contains 17 KSP motif repeats, in contrast to the 5 repeats in the *R. norvegicus*-derived constructs employed here and the 7 repeats mutated in the mouse model^30^.

In the NFH tail, by contrast, our data support that the preponderance of KSP motifs allow a conformational shift upon phosphorylation due to charge repulsion between the phosphates and the N-terminal negatively charged block. This phosphorylation-dependent expansion is in agreement with findings from Monte Carlo simulations, bovine NF hydrogels, and prior SCFT ^11,18,19,39–41^. However, the ternary NF brushes measured here were only slightly expanded by phosphorylation. This limited effect may be due to the relatively low quantity of NFH present in a 7:3:2 mixture, as the height of the binary NFL:NFH brush series seemed to correlate approximately linearly with NFH mole fraction. Previous studies in mice lacking the NFH tail domain showed that the NF spacing remained unchanged relative to wild type^17^. Our data suggest that NFH has only a minor contribution to overall brush height because of the low NFH density in the wild type^25,42^, which may explain why NFH tail deletion in mice does not affect NF-NF spacing.

The NFH tail may play roles other than regulation of brush height. Interestingly, in the NFH Δ462-638 variant we removed almost 1/3 of the total protein chain and did not significantly reduce the brush height at physiological ionic strength. Indeed, brush morphologies predicted by SCFT suggest that at physiological ionic strengths residues 462-638 contribute only to multilayer brush morphology and not to overall brush height. These data suggest a model where these C-terminal residues are brought along with the phosphorylated KSP repeat region of the sequence and contribute to the more peripheral layer of polymer density. Such a structure would be consistent with a previous proposal that the most C-terminal ∼190 residues of the *R. norvegicus* NFH tail act as an inter-filament crosslinking domain^6^, which might be most effective at the periphery of the brush layer.

The multilayered features of NFM and phosphorylated NFH brushes obtained by our SCFT are consistent with overall morphology predictions from previous studies^18,43^. We note the discrepancies between the experimental brush heights and those predicted by SCFT, which reflect aspects of the physical system not captured in the model. At low ionic strengths the model underpredicts brush height, which is attributed to the simplified treatment of a uniform dielectric environment with the dielectric constant of water. The lower dielectric constant of proteins in real brushes would amplify charge repulsion to increase brush height, and the dielectric contrast between the brush and the substrate would also increase brush height via image charge repulsion^31^. At intermediate ionic strengths, the SCFT overestimates the height of phosphorylated NFH brushes, which can be attributed to the lack of explicit treatment of proton dissociation^44^. For polyanions in this range of ionic strengths, the local proton concentration is higher than the bulk value, resulting in fewer ionized groups in the brush and therefore lower brush heights. As the ionic strength increases, sodium ions from the bulk solution displace the local hydrogen ions and the local pH approaches that of the bulk, improving the agreement between the theoretical predictions and experiments.

Our highly controlled reconstitution paradigm could incorporate additional complexities of the physiological system not readily accessible to *in vivo* studies. For example, future studies could investigate the effect of total grafting density, composition-dependent inter-brush adhesion forces, the tail domains of peripherin and α-internexin, or species-specific sequence features such as the additional KSP repeats in bovine NFM. Further, while our study focuses on phosphorylation sites in KSP repeats, future studies could explore other phosphorylation motifs recognized by non-proline-directed kinases. Finally, AFM height measurements give a label-free metric of IDP conformations in the context of a surface-tethered protein assembly. Because the brush height metric used here is based on a force resistance threshold, it may be particularly well suited to capture the biomechanical function of these brushes in resisting inter-filament compression^11,45^.

## Methods

No unexpected or unusually high safety hazards were encountered.

### Cloning

The NF protein sequences used were adapted from *Rattus norvegicus* (UniProt accession P19527, P12839, F1LRZ7) and cloned into pET28a(+) (Novagen). A cysteine was introduced near the N-terminus of each sequence to enable the grafting chemistry. For NFM and NFH tails, one tryptophan and one tyrosine residue were introduced at the N terminus to enable protein tracking during purification. NFL and NFH tail sequences include a C-terminal hexahistidine tag for purification, while NFM tail includes an N-terminal hexahistidine tag and thrombin cleavage site derived from the pET28a vector, as a C-terminal-His-tagged version of this construct was found to yield only minimal protein expression in *E. coli*. NFH Δ36-72 and NFH Δ462-638 constructs were cloned by inverse PCR from the pET28a-NFH tail plasmid.

NFM charge-shuffle sequences were designed in Python by replacing the glutamate, aspartate, residues from the NFM sequence with placeholder values, then replacing residues in a charge-alternating pattern, choosing randomly between glutamate/aspartate and arginine/lysine. Excess glutamates were placed at random among the charged residue locations. Five such sequences were randomly generated and the two with the lowest κ parameters were chosen as NFM shuffle 1 and NFM shuffle 2. DNA sequences were generated using codons optimized for *E. coli*, and genes were synthesized (Genscript), amplified by PCR, and ligated into the pET28a vector using the same restriction sites as for the wild type NFM construct. Rat ERK2 and human MKK plasmids were obtained as previously described^23^. For all amino acid sequences used in this work see Supporting Fig. 1.

### Protein production and purification

*E. coli* Rosetta (DE3) cells (Novagen) carrying a plasmid with the gene of interest were grown in Terrific Broth to an OD600 of ∼0.8-1.0, then induced with 0.4 mM isopropyl ß-D-1-thiogalactopyranoside (Thermo Fisher) according to the vector standard protocol. Protein expressions were carried out for either 4-5 h at 37 °C (NFL, NFM, NFM Shuffle 1/2, ERK2, MKK) or 16-18 h at room temperature (NFH, NFH Δ36-72, NFH Δ462-638). Cells were incubated with lysis buffer (20 mM Tris, 300 mM NaCl, 10 mM imidazole, 1 mM PMSF, 1x EDTA-free Pierce Protease Inhibitors (Thermo Scientific), ∼25 ug/mL DNAse I, ∼25 ug/mL RNAse A, ∼ 0.1 mg/mL lysozyme, pH 8.0), stirred for 2 h at 4 °C, then tip sonicated at 4 °C for 10 min total, with duty cycle 2 s on / 4 s off. Lysates were centrifuged at 20,000 xg for 30 min. For NFL, NFM, and NFM shuffles 1 and 2, proteins were purified from the insoluble fraction following lysis. These cell lysis pellets were resolubilized for several h in a denaturing buffer (8 M urea, 0.3 M NaCl, 10 mM imidazole, 20 mM Tris, pH 8.0) and centrifuged at 20,000 xg for 30 minutes, after which the supernatant containing resolubilized protein was reserved for purification. For convenience to limit buffer exchanges for these proteins, chromatography for these proteins was performed in buffers containing 8 M urea before stepwise dialysis of the purified proteins into storage buffer.

Proteins were purified by fast protein liquid chromatography (FPLC). Lysates or resolubilized protein were purified by nickel-based affinity chromatography (HisTrap HP, Cytiva) with a gradient elution from 10 to 250 mM imidazole. The NF tail constructs were exchanged into appropriate buffers and further purified by anion exchange (DEAE FF, Cytiva) in 1 mM DTT and either 20 mM MES pH 6.0 (NFL), 20 mM Tris pH 8.0 (NFM), or 20 mM Tris pH 7.5 (NFH) with a gradient elution from 0 to 2 M NaCl. For NFM, additional purification by size-exclusion chromatography (SEC) was required. SEC (Sephacryl S 300 HiPrep 16/60, Cytiva) was performed in 8 M urea, 50 mM sodium phosphate, 150 mM NaCl, 5 mM DTT, pH 7.2.

Purified proteins were extensively dialyzed into phosphate buffered saline (PBS, pH 7.4), then spin-concentrated (Amicon Ultra Centrifugal Filters) to adjust the final protein concentration to ∼1 mg/mL (NFM, NFM Shuffle 1/2, ERK2, MKK) or ∼5 mg/mL (NFL, NFH, NFH Δ36-72, NFH Δ462-638) by BCA assay (Pierce BCA Protein Assay, Thermo Scientific). Protein aliquots were stored at -80 C.

### Protein grafting

Protein was grafted to glass or silicon surfaces as described previously^20^. Briefly, glass coverslips (#48380046, VWR) or cut silicon wafers (#16010, Ted Pella) were plasma cleaned, then functionalized sequentially with 3-glycidyloxypropyl trimethooxysilane (Sigma), diamino-polyethylene glycol with molecular weight 400 Da (#PSB-3640, Creative PEGWorks), and 3-(maleimido)propionic acid N-hydroxysuccinimide ester (Sigma). For co-grafted brushes, protein and tris (2-carboxyethyl) phosphine (TCEP) stock solutions were combined to yield a constant total volume of 16 μL and final buffer composition of PBS and 1 mM TCEP. The mixture was sandwiched between two functionalized coverslips or wafer pieces and incubated at room temperature for 20 h, then washed thrice in PBS and thrice in deionized water.

### Brush phosphorylation

Protein brushes were phosphorylated by ERK2 as described previously^23^. Phosphorylated residues were confirmed by LC-MS after in-gel digestion with trypsin (Promega #V511A) according to standard protocols. Briefly, several μg of each protein were phosphorylated in solution as described previously^23^ and separated by SDS-PAGE. Gels were fixed with methanol and acetic acid and stained with Coomassie Blue. Gel bands were excised and dehydrated in acetonitrile, reduced with DTT and alkylated with acrylamide, then rehydrated with 10 ng/μL trypsin in 100 mM NH_4_HCO_3_ buffer and incubated overnight at 37 °C. Peptides were extracted in 50% acetonitrile, 5% formic acid, 45% water, and dried with a vacuum centrifuge before mass spectrometric analysis. Samples of purified intact proteins and trypsin-digested proteins were submitted to the QB3/Chemistry Mass Spectrometry Facility at the University of California, Berkeley and analyzed using Synapt and Orbitrap instruments, as described elsewhere^46,47^.

### Spectroscopic ellipsometry

Thicknesses of single-component protein brushes on silicon wafers were measured with a variable angle spectroscopic ellipsometer (M-2000V, J.A. Woollam). Samples were equilibrated in deionized water and dried thoroughly under nitrogen immediately before measurement. Spectra were taken at 50, 60, and 70° angles and were fitted as described previously^20^ using a two-layer Cauchy model over a standard silicon substrate model in the CompleteEase software (J.A. Woollam). The first layer thickness and refractive index were fitted from a one-layer Cauchy model using 2-6 reference samples included in each experiment, which were functionalized in the same batch of samples but were not exposed to any protein. Protein layer refractive indices *n* were set based on protein-sequence-dependent estimates of *n* following the method from ^48^, yielding *n* ranging from 1.585 (NFH tail) to 1.616 (NFL tail) . Grafting density was calculated as 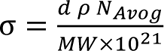, where d is the protein layer thickness, ρ is the density of a pure protein 1.41 g/mL^49^, N_Avog_ is Avogadro’s number and MW the molecular weight of each protein (g/mol).

### Fluorophore conjugation

To enable fluorescent mixed brushes, amine-reactive fluorophores chosen to minimize spectral overlap were used to label NF tail proteins. For quantifying binary brush composition, NFL was labeled with 5(6)-carboxyfluorescein NHS ester (Sigma), while NFH or NFM were labeled with Sulfo-Cyanine5 NHS ester (Lumiprobe). For a ternary brush Cyanine7.5 NHS ester (Lumiprobe) was used to label NFM. All fluorophore bioconjugation reactions were performed following manufacturer protocols (Thermo Scientific). Briefly, fluorophores were resuspended in dimethylformamide and an amount of dye corresponding approximately to an 8:1 fluorophore to protein molar ratio was combined with one of the purified proteins, prepared at 0.4 - 1 mg protein mL^-1^ in PBS adjusted to pH 8.4. Reactions were shaken at room temperature for 1-4 h, then unreacted dye was quenched for 30 min by addition of Tris (pH 8) to 50 mM. Buffer was exchanged back to PBS and the labeled protein was separated from unreacted free dye with a PD-10 desalting column (Cytiva). Labeled proteins were spin-concentrated (Amicon Ultra Centrifugal Filters) to adjust the protein concentration to match that of unlabeled protein stocks, and aliquots were frozen at -80 C. To check micron-scale homogeneity, representative images of labeled protein stock solution or fluorescent mixed brush samples were taken by widefield epifluorescence microscopy (Eclipse TE2000-E, Nikon) with Plan Fluor 20x Ph1 DLL or Plan Fluor 40x Ph2 DLL objectives.

### Fluorescent brush quantification

Fluorescent mixed brushes were formed by grafting fluorescently labeled protein onto functionalized glass coverslips. To enable quantification, each experiment included several single-component reference samples for each protein. After washing and equilibrating in PBS, the coverslips were imaged at a resolution of 25 μm (Typhoon FLA 9000, Cytiva). Samples labeled with Cyanine7.5 NHS ester were imaged with a near-infrared scanner (Odyssey, Li-Cor). Image quantitation was performed in ImageJ (Fiji). Average pixel intensity was recorded for each sample and channel, and was normalized to the average signal from the relevant single-component brush references for each channel. Assuming constant grafting density across experiments from a given protein stock, the single-component brush density is known from spectroscopic ellipsometry measurements. Assuming that signal intensity correlates linearly with labeled protein abundance, the grafting density of each component in a sample is given as 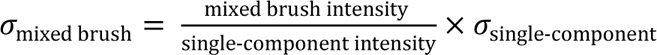. Total grafting density is found by summing grafting density over all protein components. Molar composition is calculated as 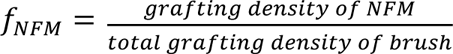 or 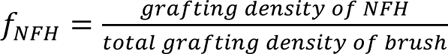. All regression fitting was performed in GraphPad Prism.

### Atomic force microscopy (AFM)

Brush height measurements were performed by AFM (MFP3D-BIO, Oxford Instruments) as previously described^20^. Briefly, AFM silicon nitride probes of nominal spring constant 0.08 N/m and nominal tip radius 20 nm (MLCT-BIO, Bruker) were calibrated by the thermal method before use and cleaned by 4 min plasma treatment. Force spectroscopy was performed under aqueous solutions at room temperature. Each force curve was 600 nm in length with a ramp rate of 1.2 μm/s and used a trigger threshold of 50 nm deflection (corresponding to ∼3 nN). To avoid re-probing the same part of the brush, replicate curves were obtained in force maps of 100 or 144 curves spaced by at least 0.2 um in both X and Y directions, and three such maps were performed at distant locations on the sample giving 300-432 replicate measurements per sample. When changing solution conditions, samples were allowed to equilibrate in the new solution for at least 90 min before measurement and the probe was rinsed with several mL deionized water and several mL of the new solution.

Raw data were imported into MATLAB for brush height extraction. The recorded Z-piezo position was converted into tip-sample distance and the pre-contact baseline region was fitted to a line and subtracted from the entire force curve. The height of the first point of contact with the brush was considered as the first point where the recorded force exceeded the baseline by three baseline standard deviations for three consecutive data points. The glass position was obtained by performing a linear fit to the furthest linear contact region, measuring standard deviation around this fit, and finding the last point at which three consecutive data points fall outside of three standard deviations of this fit. The brush height was then calculated as the difference between these two positions.

### Self-consistent field theory (SCFT)

An SCFT was applied as described previously^31^. A detailed derivation is presented in Supporting Note 1 and details on charge density and Flory-Huggins parameters are presented in Supporting Note 2. In a semicanonical ensemble, the brush proteins are modeled as multiblock charged polymers grafted upon a non-interacting substrate in a monovalent salt solution. Each protein is treated as a Gaussian chain with N number of segments of volume *v* and Kuhn length *b*. After following the standard procedures (^31^, refs. 48—52) to decouple the interacting system, and using the saddle point approximation, we obtain a series of continuous-space, self-consistent field equations. Assuming the protein brush to be homogeneous in the directions parallel to the grafting surface, the protein density profiles *ɸ_p_*(*z*) describe the distribution of residues in the direction normal to the substrate. Amino acid sequences can be mapped to the charged multiblock polymer model by grouping sections of residues which contribute nearly constant slope to the cumulative sum of the protein charge distribution. The density distribution of block *i* is related to the overall density by 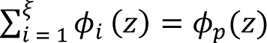 for *ξ* number of total blocks per amino acid sequence. The charge density of each amino acid is calculated at the bulk pH using the Henderson-Hasselbalch equation. The charge density *α_i_* of each block is thus the average of its constituent amino acids.. The hydrophobicity of block *i* is manifested by its Flory-Huggins interaction parameter *χ_i_*, which describes the short-range van der Waals interactions between the block and the solvent. The coarse-grained charge and hydrophobicity distributions for each protein used in the SCFT calculations are presented in Supporting Tables 2 A−H.

Pure brushes composed of NFL, NFM, NFH, and NFH deletion construct tails were modeled using the single-component brush grafting densities measured by spectroscopic ellipsometry. Reported relative hydrophobicities^50^ were rescaled such that *χ_max_* = 2.25 and the charges of phosphoserine and phosphothreonine were set at −1.0 *e*, chosen for the best fit with experimental data. For simplicity, the same Kuhn length *b* = 0.60 nm and segment volume *v* = 0.05 nm^3^ were chosen based on best fit with the experimental brush heights for all proteins except phosphorylated NFH and phosphorylated NFH deletion constructs, where *v* = 0.15 nm^3^ provided a better fit. The bulk pH was taken to be 7.44. For simplicity, the solvent was chosen to be pure water and the relative dielectric constant of the system was assumed to be uniform at the value of water, *∈_r_* = 80. The temperature was set at 293 K. To compare to brush heights measured by AFM, heights *H* were determined using a criterion where the protein density reaches a threshold value of 10^−4^.

### Net charge per residue and κ calculations

Locally averaged net charge per residue was calculated in Python for each sequence as the average charge within a moving window along the sequence length, with a window size of 31 residues. Moving average charge was not calculated for the first and last 15 residues due to smaller window size. Charge contributions were used as follows: glutamate/aspartate contributed -1.0 e, lysine/arginine contributed +1.0 e, and phosphorylated serines and threo nines at sites detected by LCMS contributed -1.5 e. κ parameters of NFM tail and NFM tail shuffle variants were calculated for sequences according to the method given in^32^ using a blob size of 5.

## Supporting Information

Supporting Information: protein sequences; validation data for protein identity and protein brush density, phosphorylation, and homogeneity; power-law exponents describing ionic strength dependence of all brush compositions; SCFT predictions of NFH Δ36-72 and NFH Δ462-638 polymer density and brush height; tabulated parameters used in SCFT calculations; and detailed derivations of the SCFT and related parameters (PDF).

## Acknowledgements

This work was supported by NIH R01GM122375 to SK and the National Science Foundation Graduate Research Fellowship under Grant No. DGE 2146752 to EAD and TJY, and used computational resources provided by the Kenneth S. Pitzer Center for Theoretical Chemistry. RW acknowledges the donors of the American Chemical Society Petroleum Research Fund for partial support of this research. We thank Dr. Anthony Iavarone of the QB3/Chemistry Mass Spectrometry Facility for mass spectrometry assistance.

