## Supplemental Information for "Dissecting neurofilament tail sequence-phosphorylation-structure relationships with multicomponent reconstituted protein brushes"

NFL tail

GCETRLSFTSVGSITSGYSQSSQVFGRSAYSGLQsSSYLMsARAFPAYYTSHVQEEQSEVEETIEATKAEEAKDEPPS
EGEAEEEEKEKEEGEEEEGAEEEEAAKDESEDAKEEEGGEGEEEDTKESEEEEKKEESAGEEQAAKKKDLEHHHHH
H

NFM tail

GSSHHHHHHSsGLVPRGSHMCWYSTFSGSITGPLYTHRQPSVtISSKIQKTKVEAPKLKVQHKFVEEIIEETKVEDEKS
EMEDALTVIAEELAASAKEEKEEAEEKEEEPEVEKsPVKsPEAKEEEEGEKEEEEEGQEEEEEEDEGVKSDQAEEGG
SEKEGSSEKDEGEQEEEGETEAEGEGEEAEAKEEKKTEGKVEEMAIKEEIKVEKPEKAKsPVPKsPVEEVKPKPEAKA
GKDEQKEEEKVEEKKEVAKEsPKEEKVEKKEEKPKDVPDKKKAESPVKEKAVEEMITITKSVKVSLEKDTKEEKPQQQ
EKVKEKAEEEGGSEEEVGDKsPQESKKEDIAINGEVEGKEEEEQETQEKGSGQEEEKGVVTNGLDVSPAEEKKGED
RSDDKVVVTKKVEKITSEGGDGATKYITKSVTVTQKVEEHEETFEEKLVSTKKVEKVTSHAIVKEVTQGD

NFH tail

GCWYMSEFTSMSTHIKVKSEEKIKVVEKSEKETVIVEEQTEEIQVTEEVTEEEDKEAQGEEEEEAEEGGEEAATTSPP
AEEAASPEKEtKSPVKEEAKsPAEAKsPAEAKsPAEAKsPAEVKsPAVAKsPAEVKsPAEVKsPAEAKsPAEAKsPAEVKS
PATVKsPGEAKsPAEAKsPAEVKsPVEAKsPAEAKsPASVKsPGEAKsPAEAKsPAEVKsPATVKsPVEAKsPAEVKsPVT
VKsPAEAKsPVEVKsPAsVKsPSEAKsPAGAKsPAEAKsPVVAKsPAEAKsPAEAKPPAEAKsPAEAKsPAEAKsPAEAKs
PAEAKsPVEVKsPEKAKsPVKEGAKSLAEAKsPEKAKsPVKEEIKPPAEVKsPEKAKsPMKEEAKsPEKAKTLDVKsPEA
KtPAKEEAKRPADIRsPEQVKsPAKEEAKsPEKEETRTEKVAPKKEEVKsPVEEVKAKEPPKKVEEEKTPAtPKTEVKES
KKDEAPKEAQKPKAEEKEPLTEKPKDSPGEAKKEEAKEKKAAAPEEETPAKLGVKEEAKPKEKAEDAKAKEPSKPSE
KEKPKKEEVPAAPEKKDTKEEKTTESKKPEEKPKMEAKAKEEDKGLPQEPSKPKTEKAEKSSSTDQKDSQPSEKAPE
DKLLEHHHHHH

NFH Δ36-72

GCWYMSEFTSMSTHIKVKSEEKIKVVEKSEKETVAATTSPPAEEAASPEKETKSPVKEEAKSPAEAKSPAEAKSPAEA
KSPAEVKSPAVAKSPAEVKSPAEVKSPAEAKSPAEAKSPAEVKSPATVKSPGEAKSPAEAKSPAEVKSPVEAKSPAE
AKSPASVKSPGEAKSPAEAKSPAEVKSPATVKSPVEAKSPAEVKSPVTVKSPAEAKSPVEVKSPASVKSPSEAKSPA
GAKSPAEAKSPVVAKSPAEAKSPAEAKPPAEAKSPAEAKSPAEAKSPAEAKSPAEAKSPVEVKSPEKAKSPVKEGAK
SLAEAKSPEKAKSPVKEEIKPPAEVKSPEKAKSPMKEEAKSPEKAKTLDVKSPEAKTPAKEEAKRPADIRSPEQVKSP
AKEEAKSPEKEETRTEKVAPKKEEVKSPVEEVKAKEPPKKVEEEKTPATPKTEVKESKKDEAPKEAQKPKAEEKEPLT
EKPKDSPGEAKKEEAKEKKAAAPEEETPAKLGVKEEAKPKEKAEDAKAKEPSKPSEKEKPKKEEVPAAPEKKDTKEE
KTTESKKPEEKPKMEAKAKEEDKGLPQEPSKPKTEKAEKSSSTDQKDSQPSEKAPEDKLLEHHHHHH

NFH Δ462-638

GCWYMSEFTSMSTHIKVKSEEKIKVVEKSEKETVIVEEQTEEIQVTEEVTEEEDKEAQGEEEEEAEEGGEEAATTSPP
AEEAASPEKETKSPVKEEAKSPAEAKSPAEAKSPAEAKSPAEVKSPAVAKSPAEVKSPAEVKSPAEAKSPAEAKSPAE
VKSPATVKSPGEAKSPAEAKSPAEVKSPVEAKSPAEAKSPASVKSPGEAKSPAEAKSPAEVKSPATVKSPVEAKSPA
EVKSPVTVKSPAEAKSPVEVKSPASVKSPSEAKSPAGAKSPAEAKSPVVAKSPAEAKSPAEAKPPAEAKSPAEAKSP
AEAKSPAEAKSPAEAKSPVEVKSPEKAKSPVKEGAKSLAEAKSPEKAKSPVKEEIKPPAEVKSPEKAKSPMKEEAKSP
EKAKTLDVKSPEAKTPAKEEAKRPADIRSPEQVKSPAKEEAKSPEKEETRTEKVAPKKEEVKSPVEEVKAKEPLEHHH
HHH

NFM Shuffle 1

GSSHHHHHHSSGLVPRGSHMCWYSTFSGSITGPLYTHKQPSVTISSEIQKTEVKAPELEVQHDFVKEIIEETRVEKEK
SEMEEALTVIAKELAASAEEKEKEAKEKDKEPKVEKSPVESPKAEKEEEGEEKDKEKGQEKEKEREKGVESKQAEEG
GSEKEGSSKEEEGKQEKEGKTDAKGEGKEAKAEKEKDTKGEVKEMAIEEKIEVEEPKDAKSPVPESPVKEVEPEPKA
DAGKEKQEKDEEVKEKDKVADESPEKEKVDEDKEKPEEVPEKEKAESPVKEKAVEEMITITESVKVSLEKDTKEEEPQ
QQEDVEEKAEKEGGSKEEVGEKSPQESEEKEIAINGEVEGKEKDKQETQEEGSGQEEKDGVVTNGLEVSPAEKEEG
EKESEEEVVVTEKVEKITSEGGKGATEYITESVTVTQEVKEHEETFKEKLVSTEKVEEVTSHAIVDKVTQGE

NFM Shuffle 2

GSSHHHHHHSSGLVPRGSHMCWYSTFSGSITGPLYTHEQPSVTISSEIQKTEVEAPELKVQHEFVKEIIEETEVDKEES
EMEEALTVIAEELAASAEEKEKEAKEKEKDPEVEKSPVESPEAEKDKEGKDEEKEKGQEKEKEKDKGVDSEQAEKGG
SEKEGSSKEKEGKQEKDGETEAEGEGKEAKAEKEKETKGEVKDMAIEEEIEVKDPKDAKSPVPDSPVKEVEPEPEAD
AGKEKQEKEREVKDKEEVAEKSPEEEKVEEEKEEPEKVPEKDKAESPVKEEAVERMITITESVKVSLEEETKEEEPQQ
QKDVKEKAEKEGGSKEKVGEKSPQESKEKEIAINGKVEGKEKDEQETQKEGSGQKDKEGVVTNGLKVSPAEKEEGE
KESKEKVVVTEKVEKITSEGGKGATEYITKSVTVTQEVEEHEETFEEKLVSTEKVEKVTSHAIVEEVTQGE

Supplementary Figure 1. All amino acid sequences used in this work. Phosphorylated residues identified by peptide mass spectrometry after in-gel trypsin digestion of phosphorylated proteins are indicated in lowercase for NFL, NFM, and NFH tails.

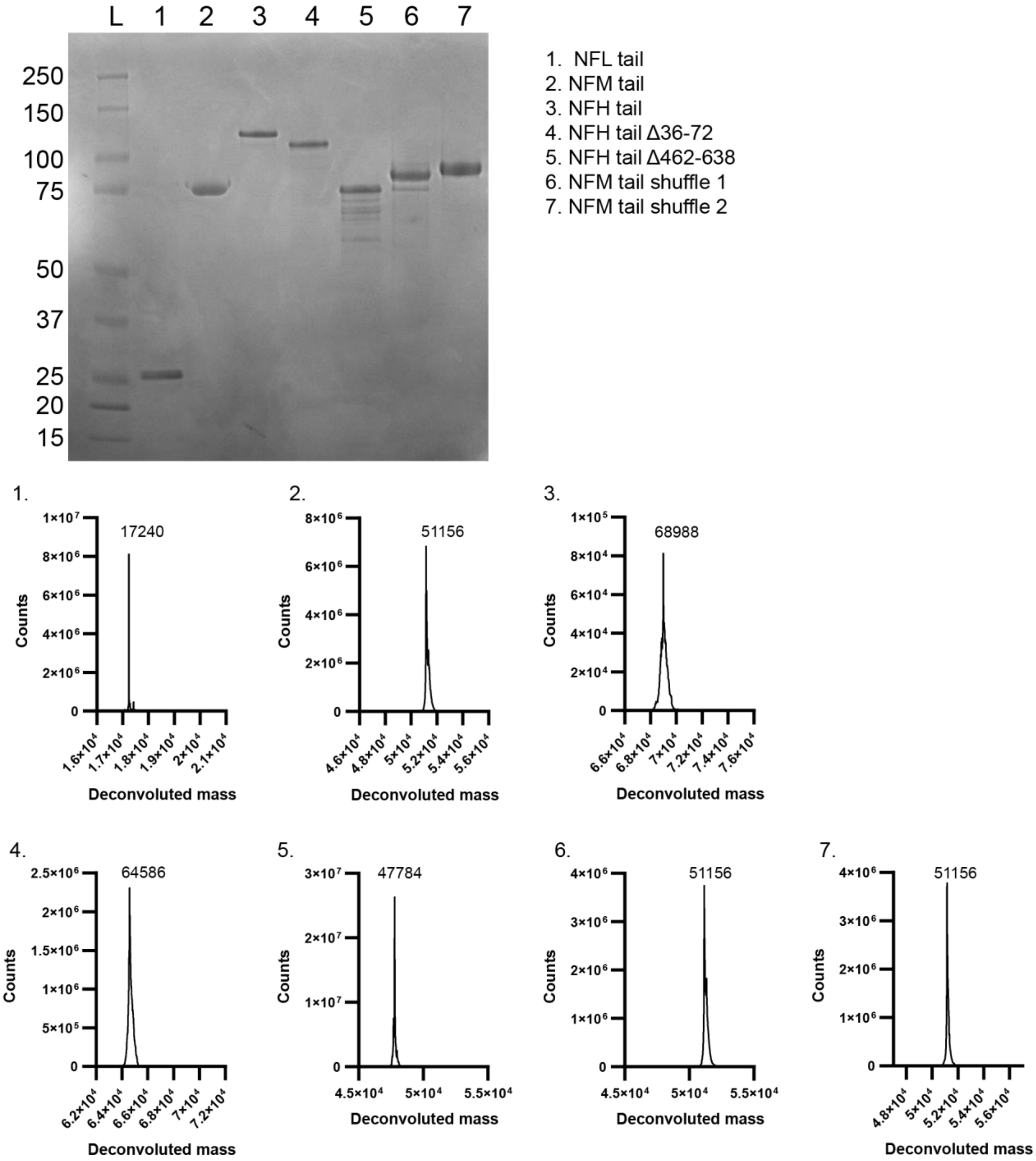


Supplementary Figure 2. (a) SDS-PAGE of all protein constructs showing anomalous migration. (b) Deconvoluted LC-MS spectra of all protein constructs. Peak masses given in Da.

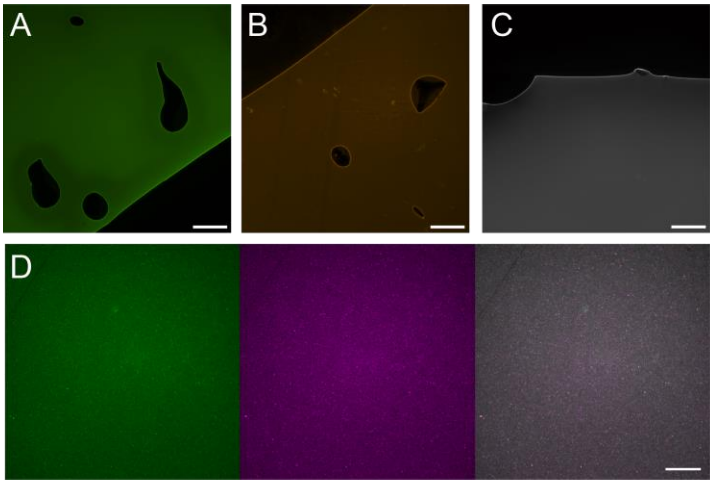


Supplementary Figure 3. (A-C) micrographs of solutions of fluorescently labeled NFL (A), NFM (B), and NFH (C) protein constructs in solution, scale bar 100 μm. Regions near bubbles are shown for contrast. (D) Representative fluorescence micrograph of mixed protein brush showing NFL labeled with FAM (green, left), NFH labeled with Cy5 (magenta, center), and composite (right), scale bar 50 μm.

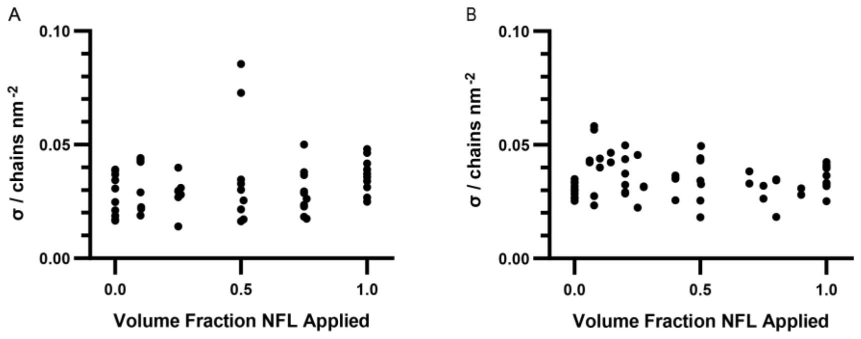


Supplementary Figure 4. Total grafting density of binary brushes as calculated from relative fluorescence measurements for NFL-NFM mixtures (A) and NFL-NFH mixtures (B). Points represent individual samples.

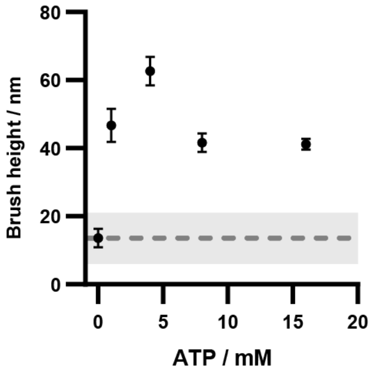


Supplementary Figure 5. Effect of ATP concentration during phosphorylation reaction for a single-component NFH brush, measured after phosphorylation at 1.6 mM ionic strength. The 0 mM ATP condition included all reaction components except ATP. 4 mM is the standard reaction condition used in this work. Points and bars represent mean and standard deviation, n=6-12. Gray line and shaded region indicate average and standard deviation brush height for a single-component NFH brush before phosphorylation reaction (n=6).

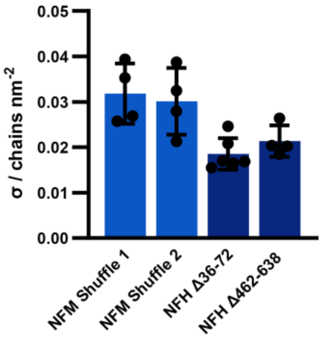


Supplementary Figure 6. Grafting densities of single-component brushes made from NFM tail shuffle and NFH tail deletion constructs. Bars show mean and standard deviation, n = 4-6.

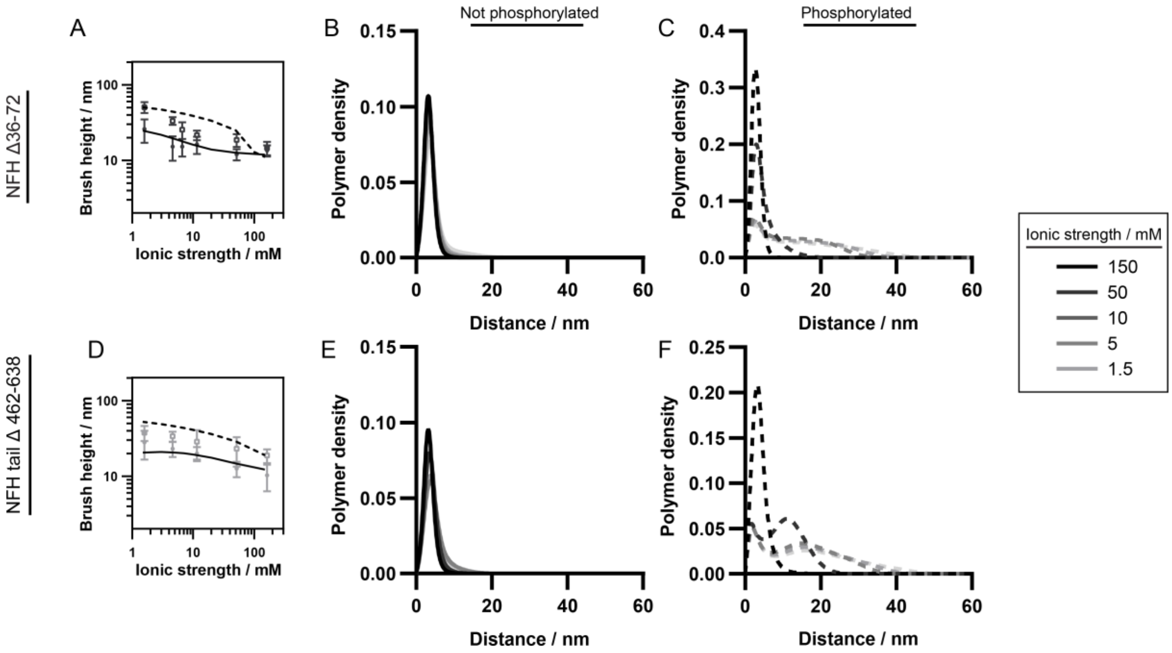


Supplementary Figure 7. Left: SCFT predicted brush heights compared to experimental brush heights for NFH Δ36-72 (A) and NFH Δ462-638 (D) constructs. Right: Predicted overall polymer density profiles at various solution ionic strengths for NFH Δ36-72 (B-C) and NFH Δ462-638 (E-F) brushes, either without (B, E) or with (C, F) phosphorylation.

| | | Not Phosphorylated | | | Phosphorylated | |
|---|---|---|---|---|---|---|
| | Stoichio-metry, $f_{NFM}$, or $f_{NFH}$ | **Regime excluded due to plateau, if any** | **Power Law Exponent** | | **Regime excluded due to plateau, if any** | **Power Law Exponent** |
| **Ternary** | 7:3:2 | | -0.4176 | | | -0.3634 |
| **NFL:NFM** | 0.28 | | -0.3261 | | | -0.2596 |
| | 0.55 | | -0.3941 | | | -0.3391 |
| | 0.77 | | -0.3591 | | | -0.2991 |
| **NFL:NFH** | 0.19 | | -0.3435 | | | -0.2337 |
| | 0.39 | | -0.3752 | | | -0.3898 |
| | 0.59 | ≥ 51.6 mM | -0.4121 | | | -0.3206 |
| | 0.79 | ≥ 51.6 mM | -0.3717 | | | -0.3580 |
| **NFL** | | | -0.2737 | | | -0.2534 |
| **NFM** | | | -0.3710 | | | -0.2856 |
| **NFH** | | | -- | | 162 mM | -0.3891 |
| **NFM Shuffle 1** | | | -0.2800 | | | -- |
| **NFM Shuffle 2** | | | -0.3021 | | | -- |
| **NFH Δ36-72** | | | -- | | ≥ 51.6 mM | -0.4401 |
| **NFH Δ462-638** | | | -0.2259 | | | -0.1521 |

Supplementary Table 1. All power law exponents. All data were fitted to the form $H = a\, c^b$, where $H$ is brush height, $c$ is solution ionic strength, $a$ is a constant and $b$ is the power law exponent reported here. Any ionic strength regimes excluded due to non-power law behavior are noted. Non-phosphorylated NFH and NFH Δ36-72 did not fit the power law model well and are excluded.

| $i$ | Residues | $\alpha_i$ | $\chi_i$ |
|---|---|---|---|
| **1** | $[0, 50]$ | $0.040014$ | $1.191774$ |
| **2** | $[51, 75]$ | $-0.278294$ | $0.822774$ |
| **3** | $[76, 125]$ | $-0.519667$ | $0.521419$ |
| **4** | $[126, 156]$ | $-0.153789$ | $0.678980$ |

Supplementary Table 2 A: NFL coarse-grained charge density and Flory—Huggins parameter distributions

| $i$ | Residues | $\alpha_i$ | $\chi_i$ |
|---|---|---|---|
| **1** | $[0, 32]$ | $0.126127$ | $1.122278$ |
| **2** | $[33, 96]$ | $-0.233632$ | $0.892061$ |
| **3** | $[97, 160]$ | $-0.437218$ | $0.556830$ |
| **4** | $[161, 192]$ | $-0.187375$ | $0.781149$ |
| **5** | $[193, 256]$ | $0.031192$ | $0.629410$ |
| **6** | $[257, 288]$ | $-0.031252$ | $0.939919$ |
| **7** | $[289, 352]$ | $-0.218616$ | $0.689289$ |
| **8** | $[353, 416]$ | $-0.077507$ | $0.890247$ |
| **9** | $[417, 438]$ | $0.047133$ | $1.021408$ |

Supplementary Table 2 B: NFM coarse-grained charge density and Flory—Huggins parameter distributions

| $i$ | Residues | $\alpha_i$ | $\chi_i$ |
|---|---|---|---|
| **1** | $[0, 35]$ | $0.001088$ | $1.101567$ |
| **2** | $[36, 70]$ | $-0.485387$ | $0.761889$ |
| **3** | $[71, 105]$ | $-0.114210$ | $0.739908$ |
| **4** | $[106, 647]$ | $0.020692$ | $0.751678$ |

Supplementary Table 2 C: NFH coarse-grained charge density and Flory—Huggins parameter distributions

| $i$ | Residues | $\alpha_i$ | $\chi_i$ |
|---|---|---|---|
| **1** | $[0, 36]$ | $0.001058$ | $1.133065$ |
| **2** | $[37, 72]$ | $-0.527420$ | $0.697984$ |
| **3** | $[73, 324]$ | $-0.130975$ | $0.823099$ |
| **4** | $[325, 647]$ | $-0.039561$ | $0.697179$ |

Supplementary Table 2 D: Phosphorylated NFH coarse-grained charge density and Flory—Huggins parameter distributions

| $i$ | Residues | $\alpha_i$ | $\chi_i$ |
|---|---|---|---|
| **1** | $[0, 35]$ | $0.001088$ | $1.101567$ |
| **2** | $[36, 67]$ | $-0.093688$ | $0.772530$ |
| **3** | $[68, 610]$ | $0.022494$ | $0.754598$ |

Supplementary Table 2 E: NFHΔ36-72 coarse-grained charge density and Flory—Huggins parameter distributions

| $i$ | Residues | $\alpha_i$ | $\chi_i$ |
|---|---|---|---|
| **1** | $[0, 35]$ | $0.001088$ | $1.101567$ |
| **2** | $[36, 286]$ | $-0.131496$ | $0.819612$ |
| **3** | $[287, 610]$ | $-0.039439$ | $0.706004$ |

Supplementary Table 2 F: Phosphorylated NFHΔ36-72 coarse-grained charge density and Flory—Huggins parameter distributions

| $i$ | Residues | $\alpha_i$ | $\chi_i$ |
|---|---|---|---|
| **1** | $[0, 35]$ | $0.001088$ | $1.101567$ |
| **2** | $[36, 70]$ | $-0.485387$ | $0.761889$ |
| **3** | $[71, 105]$ | $-0.114210$ | $0.739908$ |
| **4** | $[106, 470]$ | $0.033479$ | $0.807614$ |

Supplementary Table 2 G: NFHΔ462-638 coarse-grained charge density and Flory—Huggins parameter distributions

| $i$ | Residues | $\alpha_i$ | $\chi_i$ |
|---|---|---|---|
| **1** | $[0, 30]$ | $0.034580$ | $1.150161$ |
| **2** | $[31, 90]$ | $-0.399739$ | $0.759435$ |
| **3** | $[91, 330]$ | $-0.125032$ | $0.826694$ |
| **4** | $[331, 470]$ | $-0.062665$ | $0.767281$ |

Supplementary Table 2H: Phosphorylated NFHΔ462-638 coarse-grained charge density and Flory—Huggins parameter distributions

**Supplementary Note 1**: Derivation of Self-consistent Field Theory for Protein Brushes

In this section, a derivation of the key equations in the SCFT is provided. We have previously applied our theory to describe the morphological, scattering, and mechanical response of neurofilament-heavy brushes to varying ionic strengths and have provided a more detailed derivation [31].

For a protein brush in a semicanonical ensemble, the number of solvent molecules $n_s$ and dissolved salt ions $n_\pm$ vary to maintain the chemical potentials $\mu_s$ and $\mu_\pm$, respectively, while the number of end-tethered proteins $n_p$ remains fixed. The charged multiblock macromolecular model is used to describe proteins. Each chain with Kuhn length $b$ contains $N$ total segments and is segregated into $\xi$ number of blocks with block $i$ containing $N_i$ segments; thus, $\sum_{i=1}^{\xi} N_i = N$. The number of segments for a protein with $N_{AA}$ residues is determined by maintaining the same protein contour length and assuming an average amino acid contour length of 0.36 nm ([31], refs. 45 & 46), $Nb = 0.36\, N_{AA}$. Protein sequences are mapped onto the multiblock charged macromolecular model through a coarse-graining procedure[31], where the cumulative sum of the charge distribution is used to group similar neighboring charges into blocks. Each block is thus parameterized by the average charge density and hydrophobicity of its constituent amino acids. The procedure for obtaining amino acid charge densities and Flory—Huggins parameters can be found in Supplementary Note 2 and the resulting coarse-grained charge and hydrophobicity distributions can be found in Supplementary Table 2 A—H.

The particle-based partition function for protein brushes can be written as:

$$\Xi = \frac{1}{n_p!\, v^{Nn_p}} \prod_{\gamma} \sum_{n_\gamma=0}^{\infty} \frac{e^{\mu_\gamma n_\gamma}}{n_\gamma!\, v_\gamma^{n_\gamma}} \prod_{i=1}^{n_p} \int \mathcal{D}\{\mathbf{R}_i\} \int \prod_{j=1}^{n_\gamma} \mathrm{d}\mathbf{r}_{\gamma,j} \exp\{-\mathcal{H}\} \prod_{\boldsymbol{r}} \delta(\hat{\phi}_p(\mathbf{r}) + \hat{\phi}_s(\mathbf{r}) - 1) \ , \tag{S1}$$

where $\gamma$ denotes the small molecules in the system, $\gamma = s,\ \pm$. All possible chain configurations for protein $i$ are considered through $\int \mathcal{D}\{\boldsymbol{R}_i\}$. The final delta function enforces local incompressibility. $\hat{\phi}_p$ and $\hat{\phi}_s$ are the local instantaneous volume fractions of the polymer and solvent, respectively.

The Hamiltonian $\mathcal{H}$ in Eq. S1 includes intra-chain protein elasticity using the Gaussian chain model, short-range hydrophobic interactions between solvents and blocks, and long-range interactions between charged blocks and dissolved ions:

$$\mathcal{H} = \sum_{k=1}^{n_p} \frac{3}{2b^2} \int_0^N \mathrm{ds} \left(\frac{\partial \mathbf{R}_k(\mathrm{s})}{\partial \mathrm{s}}\right)^2 \tag{S2}$$

$$+\frac{1}{\nu}\int \mathrm{d}\mathbf{r}\ \sum_{i=1}^{\xi}\left(\frac{\chi_i}{\tilde{\nu}_p\tilde{\nu}_s}\ \hat{\phi}_i(\mathbf{r})\hat{\phi}_s(\mathbf{r})+\frac{1}{2}\sum_{j\neq i}^{\xi}\frac{\chi_{ij}}{\tilde{\nu}_p^2}\hat{\phi}_i(\mathbf{r})\hat{\phi}_j(\mathbf{r})\right)$$

$$+\frac{1}{2}\int \mathrm{d}\mathbf{r}\ \mathrm{d}\mathbf{r}'\ \hat{\rho}_c(\mathbf{r})\ C(\mathbf{r},\mathbf{r}')\ \hat{\rho}_c(\mathbf{r}')\ .$$

$\chi_i$ is the Flory—Huggins interaction parameter between block $i$ and solvent, whereas $\chi_{ij}$ is that between blocks $i$ and $j$. In this work, the cross-interactions are neglected ($\chi_{ij}=0$). $\tilde{\nu}_i=\nu_i/\nu$ is the volume of either the Kuhn segment (where $i=p$) or solvent (where $i=s$) divided by the reference volume $\nu$. Here, the reference volume is set equal to that of the solvent and taken to be water: $\nu=\ \nu_s=\nu_{water}=$ $0.030\ \mathrm{nm}^3$. The local charge density is $\hat{\rho}_c(\mathbf{r})=z_+\hat{c}_+(\mathbf{r})-z_-\hat{c}_-(\mathbf{r})+\sum_{i=1}^{\xi}\alpha_i\hat{\phi}_i(\mathbf{r})/\nu_p$, where $\hat{c}_\pm(\mathbf{r})$ is the instantaneous number of ions. $z_+$ and $z_-$ are the cation and anion valencies and $\alpha_i$ is the charge density of block $i$. $C(\mathbf{r},\mathbf{r}')$ is the Coulomb operator satisfying $-\nabla\cdot[\epsilon(\mathbf{r})\ \nabla C(\mathbf{r},\mathbf{r}')]=\delta(\mathbf{r}-\mathbf{r}')$. The scaled permittivity is $\epsilon(\mathbf{r})=kT\epsilon_0\epsilon_r(\mathbf{r})/e^2$, where $\epsilon_0$ is the vacuum permittivity, $e$ is the elementary charge, and $\epsilon_r(\mathbf{r})$ is the local dielectric constant which depends on the local composition of the system.

After following the standard self-consistent field procedure ([31], refs. 48—52) by employing particle-to-field transformations and the Hubbard—Stratonovich transformation, the resulting functional integral is replaced by the saddle-point approximation. The following self-consistent field equations are obtained after functional minimization of the free energy:

$$w_i(\mathbf{r})-w_s(\mathbf{r})=\frac{\chi_i}{\tilde{\nu}_p}\left(1-\phi_p(\mathbf{r})\right)+\frac{\alpha_i}{\tilde{\nu}_p}\psi(\mathbf{r})-\frac{\partial\epsilon(\mathbf{r})}{\partial\phi_i}\frac{|\nabla\psi(\mathbf{r})|^2}{2}\nu-\sum_{i=1}^{\xi}\frac{\chi_i}{\tilde{\nu}_p}\phi_i(\mathbf{r}) \tag{S3a}$$

$$\phi_i(\mathbf{r})=\frac{n_p}{Q_p}\int_{\sum_{j=1}^{i-1}N_j}^{\sum_{j=1}^{i}N_j}\mathrm{ds}\ q(\mathbf{r};\mathrm{s})\ q_c(\mathbf{r};\mathrm{s}) \tag{S3b}$$

$$1-\phi_p(\mathbf{r})=e^{\mu_s}\exp(-w_s(\mathbf{r})) \tag{S3c}$$

$$-\nabla\cdot\left(\epsilon(\mathbf{r})\nabla\psi(\mathbf{r})\right)=z_+c_+(\mathbf{r})-z_-c_-(\mathbf{r})+\sum_{i=1}^{\xi}\frac{\alpha_i}{\nu_p}\phi_i(\mathbf{r}) \tag{S3d}$$

$w_i$ and $w_s$ are the conjugate fields for block $i$ and solvent, respectively. $\phi_p=\ \sum_{i=1}^{\xi}\phi_i$ is the overall protein density distribution. $\psi$ is the electrostatic potential and $c_\pm=e^{\mu_\pm}/\nu_\pm\ \exp(\mp z_\pm\psi)$ is the ion concentration. $Q_p$ is the single-chain partition function, $Q_p=\ \nu_p^{-1}\int\mathrm{d}\mathbf{r}\ q(\mathbf{r};\mathrm{s})$. The propagator originating from the grafted end $q(\mathbf{r};\mathrm{s})$ in Eq. S3b satisfies the modified diffusion equation:

$$\frac{\partial}{\partial s}q(\mathbf{r};\mathrm{s})=\frac{b^2}{6}\nabla^2q(\mathbf{r};\mathrm{s})-w_i(\mathbf{r})q(\mathbf{r};\mathrm{s})\ , \tag{S4}$$

$$\text{where } w_i(\mathbf{r}) = \begin{cases} w_1(\mathbf{r}) & \text{for} \quad \mathrm{s} = [0, N_1] \\ & \vdots \\ w_\xi(\mathbf{r}) & \text{for} \quad \mathrm{s} = \left[\sum_{j=1}^{\xi-1} N_j, \sum_{j=1}^{\xi} N_j\right] \end{cases} .$$

The counter propagator originating from the free end $q_c(\mathbf{r}; \mathrm{s})$ also satisfies the modified diffusion equation.

While our theory is general for all geometries, we consider here a one-dimensional planar system, where the protein densities vary only in the $z$-direction but remain homogeneous in the $xy$-plane, to straightforwardly correspond to the experimental setup. Thus, $\sigma = n_p/A$ is the grafting density of the protein brush, where $A$ is the area of the plate. For the Poisson-Boltzmann Equation (Eq. S3d), $\mathrm{d}\psi/\mathrm{dz} = 0$ and $\psi(\mathrm{z}) = 0$ are used for the boundary conditions at $\mathrm{z} = 0$ and $\mathrm{z} = \infty$, respectively. For the modified diffusion equation (Eq. S4), $q(\mathrm{z}; \mathrm{s}) = 0$ and $q_c(\mathrm{z}; s) = 0$ are used at both boundaries. For the initial conditions, $q(\mathrm{z}; 0) = \delta(\mathrm{z} - \mathrm{z}^*)$ with $\mathrm{z}^* \to 0_+$ and $q_c(\mathrm{z}; N) = 1$.

**Supplementary Note 2**: Determination of the charge density and Flory—Huggins parameter in the charged multiblock macromolecular model

In this section, we describe how the charge density and Flory—Huggins interaction parameter are obtained for each amino acid in the protein sequence.

In this work, the charge densities of amino acids $\alpha_b^*$ are determined by their side chain dissociation constants $\mathrm{pK_a}$ and $\mathrm{pK_b}$ and the bulk pH:

$$\alpha_b^* = \frac{r_+^b}{1 + r_+^b} - \frac{r_-^b}{1 + r_-^b} \ , \quad \text{(S5)}$$

where $\log_{10} r_+^b = -(\mathrm{pH} - \mathrm{pK_b})$ and $\log_{10} r_-^b = \mathrm{pH} - \mathrm{pK_a}$ are the ratio of the number of positively and negatively charged residues, respectively, to the number of uncharged residues at bulk conditions. The charge density $\alpha_i$ of block $i$ can then be calculated by averaging the charge densities of its constituent amino acids.

The dissociation constants used in this work were reported by Lide ([31], ref. 47):

| Amino acid | $\mathbf{pK_a}$ | $\mathbf{pK_b}$ |
|---|---|---|
| D | 3.65 | – |
| E | 4.25 | – |
| H | – | 6.00 |
| K | – | 10.53 |
| R | – | 12.48 |
| C | – | 8.18 |
| Y | – | 10.07 |

The interaction parameter $\chi_i$ between block $i$ and solvent is calculated by the average of the interaction parameters of its constituent amino acids. The interaction parameters between amino acids and solvent are determined by rescaling their relative hydrophobicities[50,51]. The two endpoints, the maximum value of $\chi_{max} = \chi_F = 2.25$ and the minimum value of $\chi_{min} = \chi_D = 0.00$, were chosen for the best fit with experimental data:

| Amino acid | $\chi$ |
|---|---|
| E | 0.35 |
| K | 0.46 |
| A | 1.40 |
| P | 0.30 |
| V | 1.90 |
| T | 0.49 |
| S | 0.36 |
| D | 0.00 |
| G | 0.80 |

| Amino acid | $\chi$ |
|---|---|
| Q | 0.65 |
| H | 0.91 |
| I | 2.24 |
| M | 1.87 |
| R | 0.60 |
| F | 2.25 |
| Y | 1.71 |
| W | 2.21 |
| C | 1.51 |
